# MEF2C is a new regulator of the human articular chondrocyte phenotype

**DOI:** 10.1101/315630

**Authors:** S. Lazzarano, C.L. Murphy

## Abstract

MEF2C plays a role in diverse tissues, most notably heart, brain, eyes and developing bones. Here we report for the first time that MEF2C is present and active in the permanent articular cartilage in humans which lines and protects our joints throughout life. We show that MEF2C directly targets cartilage master regulator gene SOX9, and SOX9, in turn, regulates MEF2C in a novel positive feedback loop maintaining high expression levels of both transcription factors, and consequently stabilising the articular chondrocyte phenotype and helping prevent hypertrophy and subsequent calcification and vascularisation. We propose that MEF2C and SOX9 may show similar cooperative activity in other tissues, and across a range of adult murine tissues we found co-expression of both transcription factors in cartilage, trachea, brain, eyes and heart. Strikingly, all of these tissues are prone to calcification and further study of MEF2C/SOX9 cooperativity in these organs will be revealing.

## INTRODUCTION

Human articular chondrocytes (HACs) are the only cell population present in articular cartilage and they produce and maintain its extracellular matrix (ECM) in extreme physiological conditions such as high mechanical stresses and low levels of oxygen (hypoxia) [1-5] [6]. Intriguingly, the transcriptional apparatus that sustains the expression of the cartilage matrix is induced by hypoxia through HIF-2α targeting of cartilage master regulator transcription factor SOX9 [7-9]. Furthermore, it has been possible to identify potentially novel genes important for chondrocyte function by their up-regulation in hypoxia and down-regulation upon de-differentiation [7]). By this means we identified Myocyte enhancing factor 2 isoform C (MEF2C) as a potential gene of interest in HACs.

Studies have shown MEF2 expression in lymphocytes, smooth muscle, neural crest, endothelium, and bone with the highest levels in skeletal muscle and brain where they play a central role in specific cellular genetic programmes and stress responses [10-17]. The C isoform (MEF2C) of this family of transcription factors has been found to be crucial for bone development in mice [11]. The inducible Cre/loxP deletion of the *Mef2c* gene (by using a *Col2a1* gene promoter in a murine model) prevents chondrocyte hypertrophy during the last phases of the endochondral ossification process (mice carrying a transcriptional repressor fused with the Mef2c gene give the same phenotype). Conversely, the expression of Mef2c-VP16 fusion protein in a murine model results in premature ossification at E 8.5, which gives a phenotype with excessive ossification of the ribs and sternum, bone shortening and malformation caused by a premature ossification. This function is partially due to MEF2C regulation of type X collagen (in conjunction with SOX9), an important component of the hypertrophic ECM. Outside of bone development, the link between MEF2C and ECM biology is less clear, but Lin and colleagues highlighted that MEF2C-null embryos fail to form endocardial chambers due to inadequate ECM production [14].

We present the first evidence for a key role for MEF2C in maintenance of the permanent articular cartilage in humans. In contrast to the growth plate, HACs resist hypertrophy and subsequent calcification and vascularisation partly through maintenance of SOX9 expression [18]. In the present study we uncover a new positive regulatory loop involving MEF2C and SOX9 which crucially maintains high levels of SOX9 thus helping to stabilise the HAC phenotype, and enable the articular cartilage to resist calcification. In addition to articular cartilage we show MEF2C and SOX9 co-expression in brain, eyes, trachea and heart where we propose similar co-operative activity may occur.

Possible implications for this in these calcification-susceptible tissues are discussed.

## RESULTS

### The MEF2C mRNA and protein are induced by hypoxia in HACs

MEF2C exerts a pivotal role in the control of diverse cell differentiation programmes. However, the expression of the MEF2C has never before been related with hypoxia in adult human tissues and cells. To understand whether the expression of MEF2C protein and mRNA both can be increased by hypoxia, HACs have been cultured for 48 hours in hypoxia (1% O_2_) and normoxia (20% O_2_). In Western Blotting, MEF2C protein is visualised as a multi-band staining strongly enhanced by hypoxia (Figure 1A) that follows the up-regulation patterns of its mRNA and other chondrocyte-specific mRNAs SOX9 and COL2A1 (Figure 1B).

**FIGURE 1.**
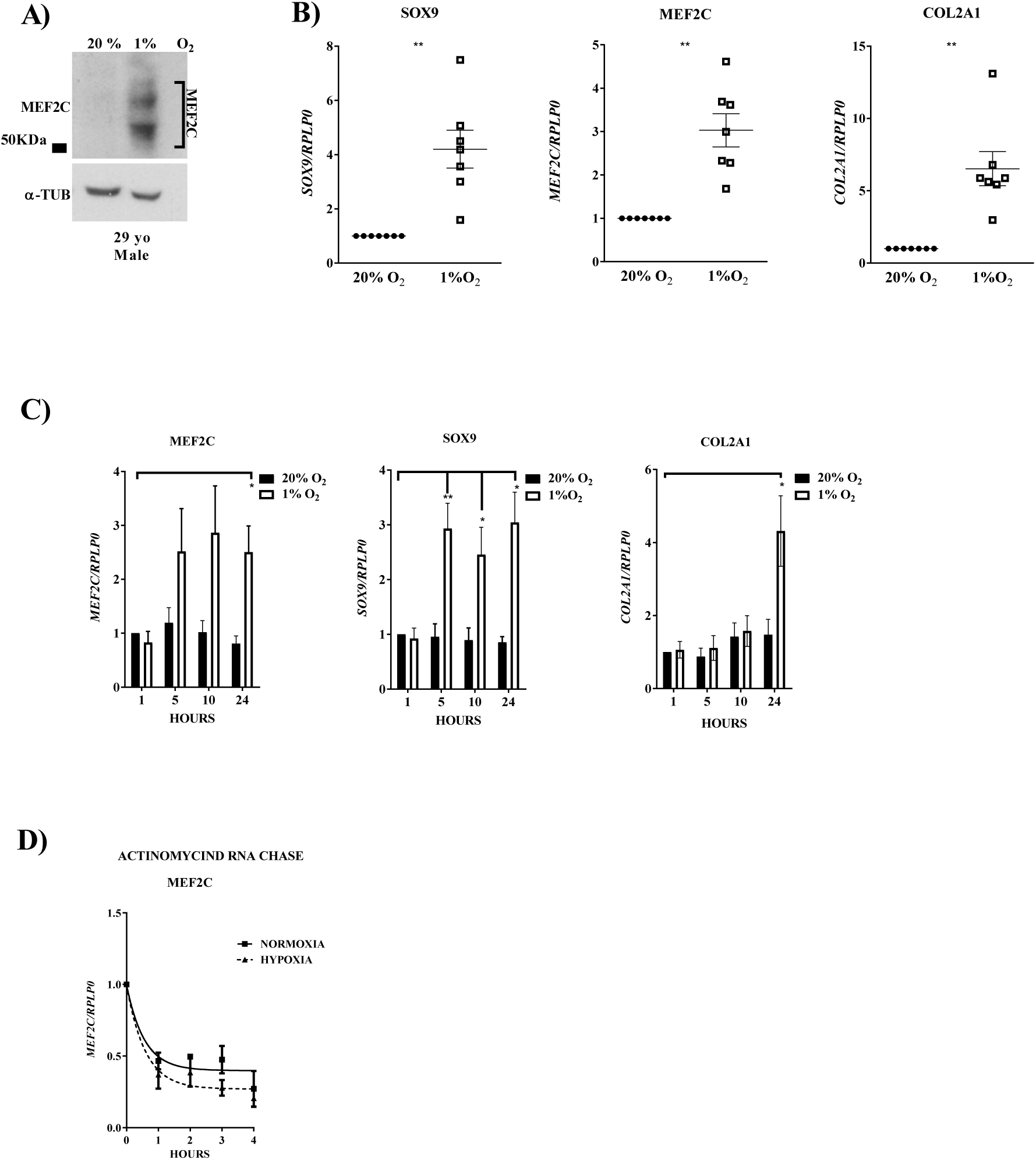
MEF2C is regulated by hypoxia in human articular chondrocytes. Western Blot showing an increase in MEF2C protein level in human articular chondrocytes (HACs) after 2 days in hypoxia (A); note: TUB – α-tubulin. SOX9, MEF2C and COL2A1 are all induced by hypoxia at the mRNA level following 2 days of treatment (B, data show the average from 7 independent experiments). HACs (passage 1 or 2 monolayer cultures) were incubated for 1, 5, 10 and 24 hours in 20% or 1% O2 and SOX9, MEF2C, COL2A were quantified using RT-qPCR and expressed relative to 1h levels in 20% O2 (C, data show the average from 4 independent experiments). Hypoxia does not affect MEF2C mRNA stability in HACs (D). HAC monolayer cultures were maintained for 1 day in normoxia (20% O2) or hypoxia (1% O2) and samples were harvested after 1, 2, 3 and 4 hours following administration of Actinomycin D (data show the average from 4 independent experiments).

In order to further characterise the regulation MEF2C gene expression, the kinetics of the MEF2C mRNA up-regulation in hypoxia was studied. In HACs cultured for 1, 5, 10, 24 hours, MEF2C mRNA shows increased transcript levels (on average) by 5 hours of hypoxia stimulation, becoming statistically significant by 24 hours (Figure 1C). These experiments show that cartilage master regulator transcription factor SOX9 is an “early response” gene and MEF2C shows a similar pattern of expression in response to hypoxia. As expected SOX9 target and key cartilage matrix gene COL2A1 shows a slower response to hypoxia. Furthermore, hypoxia did not significantly affect MEF2C mRNA stability as assessed by Actinomycin-D RNA chase assays (Figure 1D) indicating the increased mRNA levels observed are due to an increased rate of transcription.

Immunostaining of freshly isolated healthy articular cartilage from adult human knee joints showed a predominately nuclear localisation of MEF2C protein similarly to HIF-2α and SOX9 (Figure 2 A,B). Across a range of adult murine tissues (RNA from 6 mice was pooled together), gene expression of both Mef2c and Sox9 was investigated, and significant co-expression was observed in articular cartilage, trachea, brain, eyes and to a lesser extent heart (Figure 2C).

**FIGURE 2.**
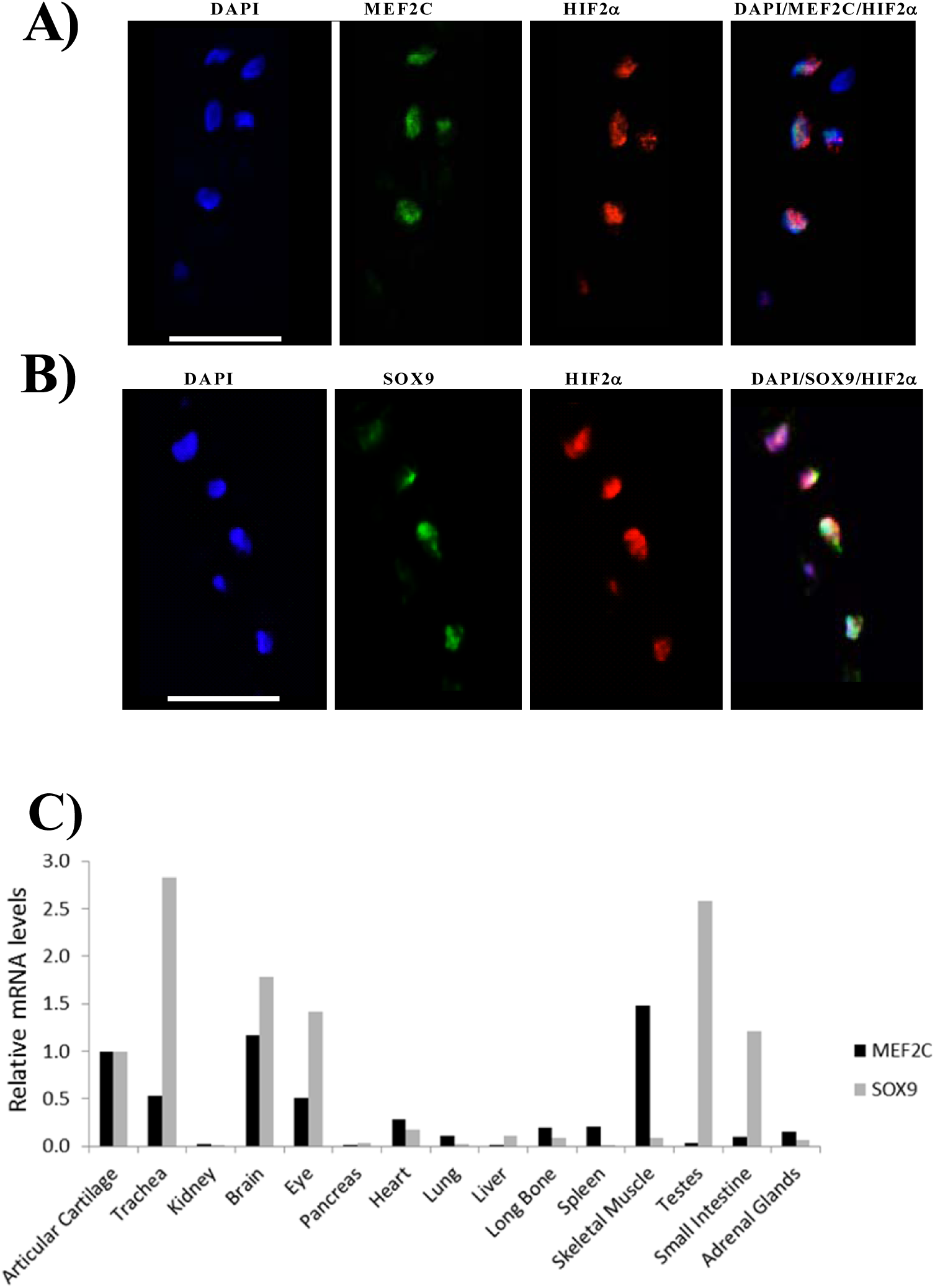
MEF2C protein is present in the nucleus of chondrocytes in intact human articular cartilage. Cryosections of freshly harvested human articular cartilage were stained for MEF2C, HIF-2α and SOX9 proteins. Nuclear co-localisation of MEF2C and HIF-2α (A) and of SOX9 and HIF-2α (B). Mef2c and Sox9 gene expression across 15 adult murine tissues (pooled samples from 6 mice). Messenger RNA levels were normalized to those of 28S and expressed relative to articular cartilage levels. Note: α-TUB – α-tubulin.

### Regulation of MEF2C expression in HACs

We next investigated the role of HIF-2α, and its main target in HACs, SOX9, on MEF2C gene expression. Following siRNA-mediated silencing of HIF-2α (encoded by the EPAS1 gene) or SOX9, HACs were cultured for two days at 20% and 1% O_2_. Interestingly, silencing of each individual transcription factor caused a decrease in MEF2C expression at both gene and protein levels (HIF-2α, Figure 3 A-B, SOX9, Figure 3 C,D). The depletion of SOX9 seemed to have a stronger effect on MEF2C levels (Figure 3 C,D). In contrast to HIF-2α, HIF-1α depletion had no impact on MEF2C expression levels (supplementary Figure 1 A).

**FIGURE 3.**
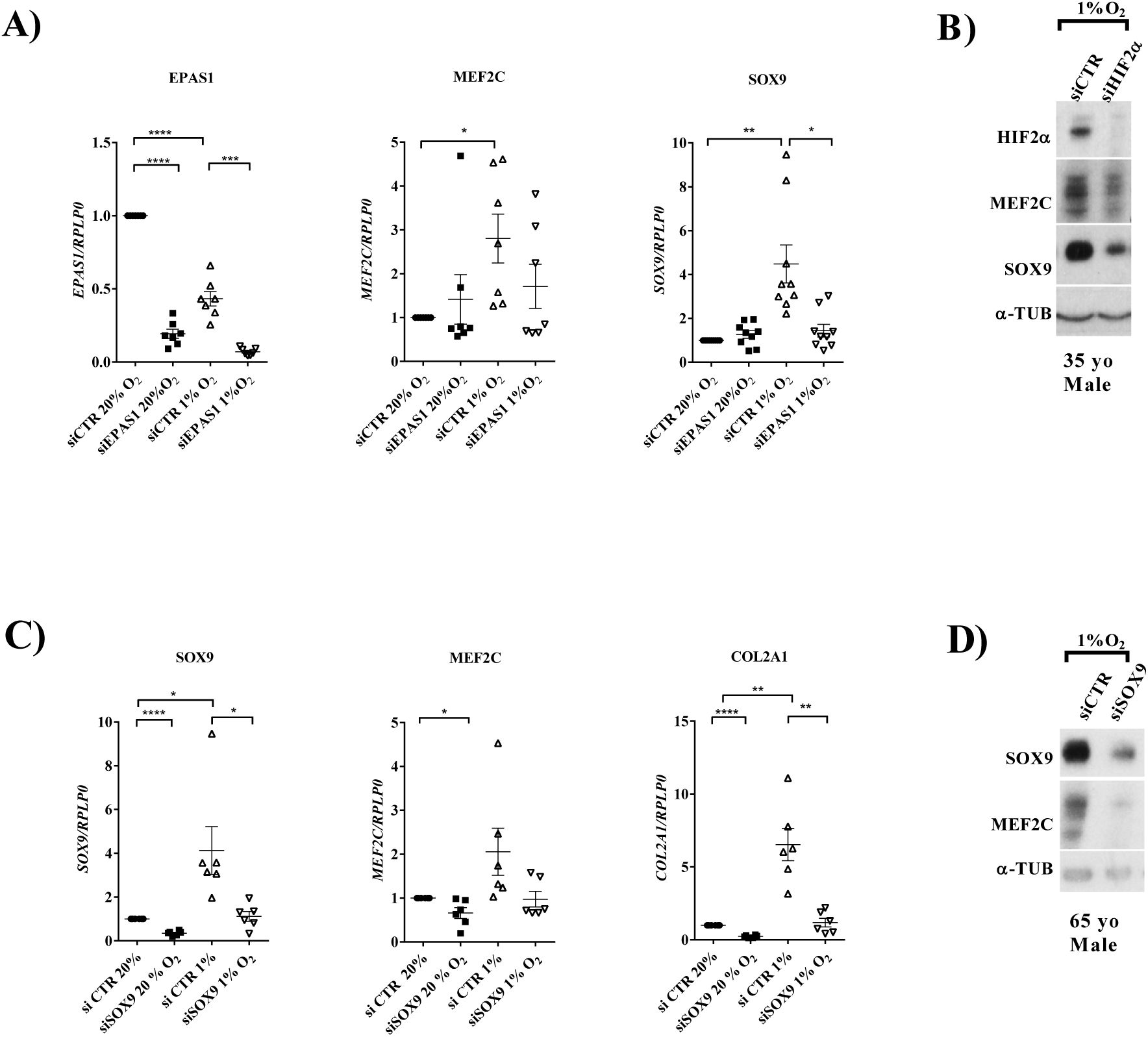
MEF2C expression is regulated by HIF-2α and SOX9 in human articular chondrocytes. Effect of depletion of HIF-2α (encoded by EPAS1) in HAC cultures results on MEF2C and SOX9 mRBA (A) and protein (B) levels. Effect of depletion of SOX9 on MEF2C and COL2A1 mRNA levels and MEF2C protein levels (D). (PCR expression data from 7 independent experiments). Note: α-TUB – α-tubulin, yo – years old; CTR – control. For display purposes blots have been cropped.

### MEF2C protein is required for SOX9 gene expression

The role of the MEF2C in HACs is unknown. In order to uncover this function MEF2C was depleted by transfection of siRNA in HACs (11 donors) which were then cultured for two days in 20% and 1% O_2_. SOX9 expression was dramatically impacted by MEF2C depletion (Figure 4 A,B); as was cartilage matrix gene and SOX9 target, COL2A1 (Figure 4A) in both normoxic and hypoxic conditions. The silencing of MEF2C in HACs has no appreciable effect on HIF-2α protein levels in hypoxia (Figure 4C). MEF2C silencing resulted in the down-regulation of another transcription factor - DLX5 (supplementary figure 2). The distal-less-homebox 5 is involved in the murine cranio-facial development along with MEF2C and has been shown to be a hypoxia inducible gene in HACs [17].However, DLX5 silencing did not alter SOX9 expression (supplementary figure 3).

**FIGURE 4.**
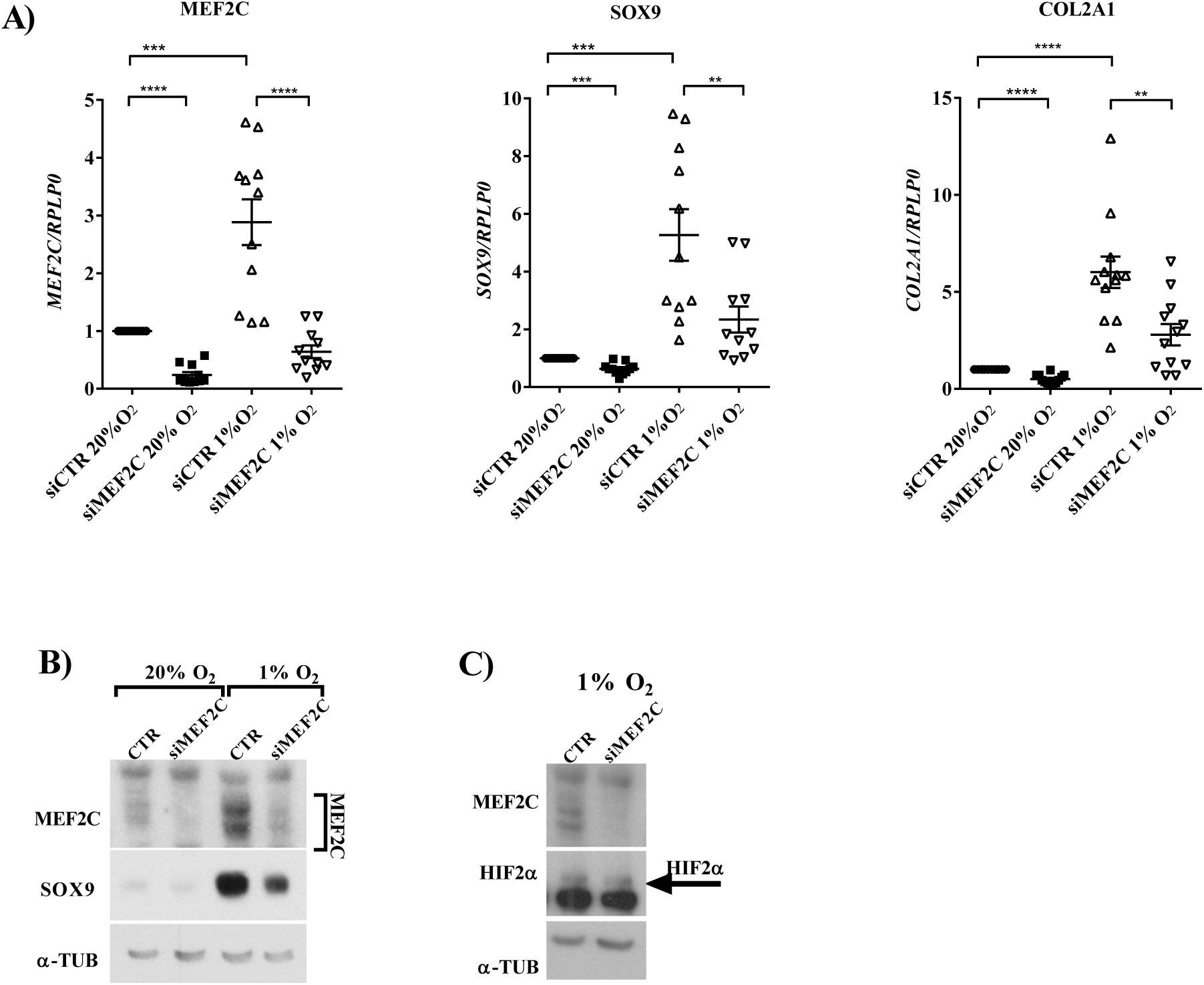
Expression of cartilage master regulator SOX9 is strongly dependent on transcription factor MEF2C in human articular chondrocytes. Following transfection with siMEF2C, HACs were cultured for 2 days in 20% or 1% O_2._ Depletion of MEF2C resulted in a significant decrease in SOX9 gene expression and of SOX9 key target gene, COL2A1 (A). A decrease in SOX9 protein levels was particularly evident in hypoxia (1% O_2_) (B), while HIF-2α levels were unaffected (C) in MEF2C depleted HACs. For display purposes blots have been cropped.

### MEF2C is directly involved in the transcriptional activation of SOX9

Having established that SOX9 is an early response gene during the HAC response to hypoxia (Figure 1C), we sought to investigate whether MEF2C transcription factor is required for the activation of SOX9 transcription. To this end we performed time course experiments in HACs exposed to hypoxia where MEF2C expression was first silenced. MEF2C depletion abolished hypoxic induction of SOX9 at mRNA and protein levels (Figure 5, A,C).

**FIGURE 5.**
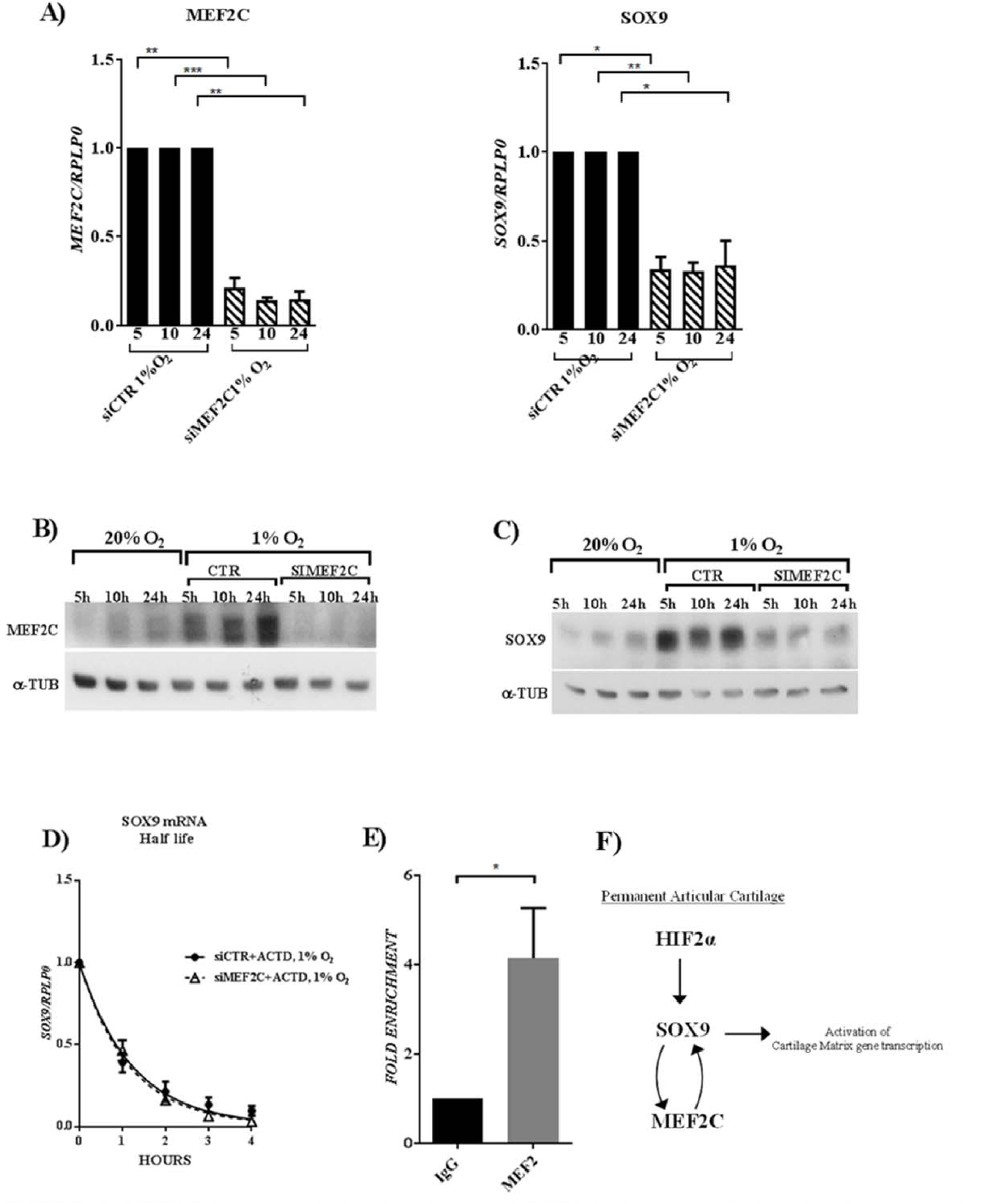
MEF2C is a trascriptional activator of SOX9 in human articular chondrocytes and is required for its hypoxic induction. The silencng of MEF2C abolished the rapid hypoxic induction of SOX9 at mRNA (A) and protein (C) levels (data from 3 independent experiments). MEF2C does not affect SOX9 mRNA stability as a revealed by actinomysin D chase experiments (D, data from 4 independent experiments). ChIP assays reveal that MEF2C directly binds to the SOX9 gene at an upsteam site (E). Schematic summarising the identified role of transcription factor MEF2C in the permanent articular cartilage in humans (F). For display purposes blots have been cropped.

In order to understand the mechanism behind the regulation of SOX9 gene expression mediated by MEF2C, we next investigated whether MEF2C can affect SOX9 mRNA stability. After transfection with specific siRNA against MEF2C, cells were cultured for 48 hours in 1% O_2_ before treatment with Actinomycin-D and harvested at 1h, 2h, 3h, and 4h. MEF2C depletion had no effect on SOX9 mRNA stability (Figure 5 D). Collectively these data strongly suggest MEF2C is (directly or indirectly) activating transcription of SOX9 mRNA.

The four MEF2 family members (A, B, C, D) have a high homology in the amino-terminal domain that regulates DNA binding and the communal minimal consensus sequence of the MEF2 transcription factor is 5′-CT(A/t)(a/t)AAATAG-3′, in tissues including skeletal muscle, cardiac muscle and brain (Andres, Cervera, & Mahdavi, 1995). We used the open-source software JASPAR (http://jaspar.genereg.net) for identifying potential binding sequences within SOX9 gene itself, as well as a 5kb upstream region (chromosome 17q24.3, the region tested was 7 kb long). As a result, we obtained a list of possible sites, which contained the 5′-CT(A/t)(a/t)AAATAG-3′ sequence. However, none of the putative sites indicated by the software turned out to be conserved between species when analysed with rVISTA software (http://rvista.dcode.org). Conservation of binding sites across different species could be an important functional indication of the involvement of that sequence into the regulation of a target gene. We have to also consider that the control of gene expression could be dramatically different among tissues/organs or different species (Wasserman & Sandelin, 2004).

Finally, we tested whether MEF2C directly binds the SOX9 gene using a Chromatin Immunoprecipitation assay (ChIP). An antibody against MEF2 previously used to study the MEF2A-C genetic network (Kalsotra et al., 2014), was used to isolate putative binding site sequences within the SOX9 gene and 5kb upstream region (chromosome 17q24.3). Our experiments show an enrichment of specific binding on the sequence situated between −4,412 bp and - 4,400 bp from SOX9 gene (Figure 5E).

## DISCUSSION

In the present study using human tissue we identify an important new role for MEF2C in the permanent articular cartilage lining our joints. SOX9 is essential for cartilage formation and function [19] and mutations in this gene cause severe skeletal defects [20].One of the key factors maintaining the permanent articular chondrocyte phenotype, and preventing calcification is maintenance of sufficiently high levels of SOX9 [18].Here we show for the first time that transcription factor MEF2C promotes SOX9 expression in human articular chondrocytes by directly targeting this cartilage master-regulator gene. SOX9 in turn positively regulates MEF2C expression in a positive feedback loop, which helps maintain high expression of both transcription factors in the permanent articular cartilage (Figure 6). We propose similar MEF2C/SOX9 co-operative function occurs in other tissues, namely heart, trachea, brain and eyes.

Articular chondrocytes maintain their function throughout life by retaining a stable phenotype that does not undergo hypertrophy and resists calcification and vascularisation. The articular phenotype is thus distinguished from the chondrocytes in the growth plate, which proliferate and undergo hypertrophy and apoptosis followed by calcification, vascularisation and bone formation. A major unresolved question in cartilage biology is what maintains the permanent articular cartilage throughout the lifespan of the organism. Maintenance of SOX9 levels seems key as this transcription factor not only regulates cartilage matrix production [7], it also acts as an important negative regulator of cartilage calcification, vascularization and endochondral ossification [18]. We previously identified hypoxia as a positive regulator of the HAC phenotype specifically through upregulation of SOX9 via binding of transcription factor HIF-2α [8]. Here we identify MEF2C as a new regulator of SOX9 in HACs. We found that MEF2C binds to a 4kb upstream site from the SOX9 proximal promoter. MEF2C expression levels were in turn positively regulated by SOX9 and it remains to be seen whether this regulation is also through direct binding. The positive feedback loop maintaining levels of both MEF2C and SOX9 suggests MEF2C may have additional important roles in HACs (other than directly regulating SOX9 levels). This will be a fruitful area of future research.

Interestingly, the role of MEF2C in the developing skeleton, where it positively regulates chondrocyte hypertrophy in the growth plate, [21]appears very different from that in the permanent cartilage, which resists hypertrophy and calcification throughout life. This highlights the fact that specific adaptations have occurred in the permanent cartilage which maintain the articular phenotype, as opposed to the growth plate where chondrocytes terminally differentiate, undergo apoptosis and are replaced by bone. Here we show that MEF2C is a critical component of this adaption in articular cartilage through its positive regulatory loop with SOX9. Interestingly in osteoarthritis - the most common joint disease - articular chondrocytes can show an altered phenotype which recapitulates some events seen in developing bone such as cellular hypertrophy, with subsequent vascularization and focal calcification [22].Therefore an important area for further investigation is the possible role of MEF2C (and changes in its expression level) in development of joint diseases such as osteoarthritis.

In addition to cartilage, across a range of murine tissues we found co-expression of Mef2c and Sox9 specifically in brain, trachea, eyes and heart. Like cartilage, these tissues can undergo calcification and it will be most interesting to further investigate Mef2c/Sox9 co-operative function in these tissues. Sox9 is critical for heart valve formation and reduced Sox9 function has been shown to lead to heart valve calcification – a problem of real clinical significance [23]. Intriguingly, Mef2C has also been show to regulate ECM production and organization in developing heart valves through direct targeting of cartilage link protein (Ctrl1)[24]. Furthermore, using chromatin immunoprecipitation assays in embryonic heart tissue the authors showed that both Mef2c and Sox9 bind to the Ctrl1 promoter. In genome-wide array studies we previously showed that MEF2C tightly clustered with CTRL1 (also known as HAPLN1) and other established cartilage matrix genes (and SOX9 targets) COL2A1, AGC and COL27A1 [7]. Thus there are striking parallels in the regulatory mechanisms between heart valves and cartilage, and further work is needed to unearth the role of MEF2C/SOX9 co-operative function in the pathophysiology of these tissues.

Krishnan et al show that HIF-1α transcription factor drives the expression of MEF2C in the heart [25]. However, we have found that MEF2C expression is specifically HIF-2α dependent in adult HACs. Whether this HIF-2α -dependence of MEF2C is cartilage-specific, or is also relevant for other hypoxic tissues that express MEF2C remains to be seen. It certainly makes sense from a cartilage perspective since we have shown that HIF-2α is a pro-anabolic regulator of human cartilage through direct regulation of SOX9 [8, 9]. In the current study it may be that the HIF-2α dependence of MEF2C is actually due to HIF-2α mediated induction of SOX9, rather than direct targeting of MEF2C by HIFs. It is of interest to note that many of the tissues co-expressing Mef2c and Sox9 are avascular/ or poorly vascularized – articular cartilage, trachea, heart valves, eyes (retina). It is interesting to speculate that MEF2C/SOX9 may be playing a role in inhibiting vascular invasion in these tissues. In cartilage SOX9 positively regulates anti-angiogenic factors such as chondromodulin [26] while *Vegfa* was shown to be negatively regulated by direct binding of Sox9 to the *Vegfa* promoter [18]. In endothelial cell-specific MEF2C-deficient mice it was shown that MEF2C appears to play an anti-angiogenic role in the retina under stress conditions, and it was suggested that modulation of MEF2C may prevent pathologic retinal neovascularization [27]. Given Sox9’s involvement in development of this tissue [28], investigation of MEF2C/SOX9 co-operative function in the retina is an exciting area for future exploration.

We conclude that MEF2C is a key new player in promotion and maintenance of the permanent articular chondrocyte phenotype. MEF2C directly regulates SOX9 expression, and that SOX9, in turn, also regulates MEF2C in a positive feedback loop which helps maintain high expression levels of both transcription factors, thereby stabilising the articular chondrocyte phenotype in the permanent cartilage lining our bones. We propose that such MEF2C/SOX9 co-operative function is important in other tissues where they are co-expressed, namely brain, trachea, eyes and heart, and this represents an exciting opportunity for future research.

## MATERIALS AND METHODS

### Human cartilage tissue

Healthy human articular cartilage was obtained from the femoral condyle and tibial plateau from amputations due to soft tissue sarcomas not involving the joint. Tissue samples were obtained after institutional approval of experiments (isolation of cells or histological processing), informed written consent, and adherence to Helsinki Guidelines (London - Riverside Research Ethics Committee, reference number 07/H0706/81). Cartilage was collected on the day of surgery and cut into small pieces before either digestion to obtain isolated cells or use for histology.

### Human articular chondrocytes (HACs)

Donor age ranged from 8 to 62 years old, including both males and females. Cartilage was collected on the day of surgery and cut into 1-2mm edged cubes. Next, the extracellular matrix proteins were digested using collagenase as previously described [8]. Cells were seeded at a density of 8×10^3^ cells/cm^2^ and cultured for 5-7 days before passaging. In the studies performed both primary (P0) and passaged cells (up to P3) were used.

### Time course experiments

The day preceding the experiment cells were counted and seeded 5 × 10^3^ cells/cm^2^ in 3.5 cm dishes. HACs (P1-P3) were exposed for 1h, 5h, 10h, and 24h to hypoxia using pre-equilibrated medium (10% FBS, DMEM). All the experiments have been performed in InVIVO_400_ Workstation (Ruskinn Technologies).

### mRNA stability assay

HACs were seeded 5 × 10^3^ cells/cm^2^ in 3.5 cm and incubated in hypoxia (1% O_2_) or normoxia (20% O_2_). After 48h hours the media were switched to media containing 5 μg/mL actinomycin D (Sigma Aldrich Biochemicals) to inhibit transcription; each hour after that, total RNA was harvested using TRIzol (Invitrogen) following the manufacturer’s protocol. The mRNA was quantify using RT-Q-PCR (standard-curve method) and half-life was calculated via GraphPad 6.0 software ( *one-decay phase equation*). All experiments in hypoxia were performed in InVIVO_400_ Workstation (Ruskinn Technologies).

### Immunofluorescence staining

Following fixation, cryosections or monolayer cultures were blocked with (5% wt/vol BSA, 1% goat serum wt/vol in 0.01% Tween/PBS) for 1hour at room temperature, and then incubated for 16 hours with primary antibody and for 1hour with appropriate secondary antibody. Cells were counterstained with 300 nM 4′-6-diamidino-2-phenylindole (DAPI) for 5 mins which was contained in the mounting agent ProLong® Antifade (Invitrogen). Antibodies used: 1:1000 SOX9 (Millipore, Ab-5535), 1:100 HIF-2⟨ (Santa Cruz, sc-13596), 1:200 MEF2C (Cell Signalling Technologies, D80C110) 1:1000 α-Tubulin (Sigma Aldrich, ab7291). High resolution digital images were acquired using a Ultraview confocal microscope (PerkinElmer) and merged using Volocity® 3D Image software ( PerkinElmer).

### Small Interfering RNA Transfection

The day preceding transfection cells were counted and seeded 5 × 10^3^ cells/cm^2^ in 3.5 cm dishes. The cells were transfected with siRNA against SOX9 (MWG ACAGAAUUGUGUUAUGUGAdTdT), MEF2C (s8653, s8654, #143535, Life technologies) HIF1α (Dharmacon GCAGUAGGAAUUGGAACATT), HIF-2α (encoded by EPAS1) (Dharmacon CGACAGCUGGAGUAUGAATT) or with negative control sequences (Silencer ® Negative Control #1 using Lipofectamine^™^ 2000 (Invitrogen). All siRNA were used at a final concentration of 10nM. After 4 hours of incubation in 20% oxygen, OptiMEM-I was replaced by previously equilibrated medium (10% FBS, DMEM). Subsequently cells were incubated at the appropriate oxygen tension (1% or 20%) for the duration of the experiment.

### RNA Extraction, Reverse Transcription, and Real Time PCR

RNA was isolated using TRIzol (Invitrogen) with an improvement to the single-step RNA isolation method developed by Chomczynski and Sacchi (Moniot, Biau et al. 2004). The concentration and quality of isolated total RNA were measured using the Nano-drop (ThermoScientific). For the murine studies total RNA from 15 different mouse tissues (pooled samples from a total of six male C57 BL/6 mice, aged 12–16 weeks) was obtained from Zyagen (San Diego, CA). Complementary DNA (cDNA) was generated using Applied Biosystem’s High capacity cDNA Reverse transcription kit with RNAse inhibitors. The reaction was prepared using 500 ug of total RNA in 20 μl of total volume and according to the manufacturer’s instructions. Newly synthesized cDNA was diluted 5 fold in DNase-free water.

For PCR, 4% of the cDNA which was reverse transcribed from 500 ng of total RNA was used. Quantitative Real Time PCR (Fast Sybr-Green Master Mix, Applied Biosystems) was performed to determine the relative expression of SOX9 (FW 5`-CGCCATCTTCAAGGCGCTGC-3′, RV 5’- CCTGGGATTGCCCCGAGTGC- 3′) MEF2C (FW 5`- ACCTATTGCCACTGGCTCAC-3`, RV 5`-ACCCATCAGACCACCTGTGT-3`) EPAS1 (HIF-2⟨) (FW 5`- AGATGGCCACATGATCTTTCTGT-3`,RV 5`-CCTGTTAGCTCCACCTGTGTAAGTC-3`), CAIX (FW 5`- TCGGAGCACACTGTGGAAG-3`, RV 5`-AAGGCCTCGTCAACTCTGG-3`) COL2A1 (FW 5`- GGAAGAGTGGAGACTACTGGATTGAC - 3′, RV 5′- TCCATGTTGCAGAAAACCTTCA - 3′) and housekeeping gene RPLP0 (FW 5`-CCATTGAAATCCTGAGTGATGTG-3`, RV 5`-CTTCGCTGGCTCCCACTTT-3`). The qPCR reactions were performed in triplicate using SYBR Green PCR Master Mix (Applied Biosystem) with appropriate primers for each gene. The standard curve method was used to calculate relative levels for each transcript.

### Western Blotting

HACs were cultured as monolayers in 20 or 1% oxygen tension for 36–48 h before lysis in radio-immunoprecipitation assay buffer (150 mM NaCl, 10 mM Tris, pH 7.2, 5 mM EDTA, 1% Triton X-100, 0.1% SDS, 1% deoxycholic acid). Proteins were detected by Western blotting using a polyclonal anti-SOX9 antibody (1:1000, AB5809 Millipore), anti-HIF-2⟨ (1:250, sc-13596,Santa Cruz), anti-MEF2C (1:500, D80C110,Cell Signalling Technologies,), α-Tubulin (1:10000, ab7291, Sigma Aldrich). and ECL reagent (Amersham Biosciences, Buckinghamshire, UK) according to the manufacturer’s instructions.

### ChIP Assays

MEF2-ChIP was performed using the Imprint chromatin immunoprecipitation kit (Sigma-Aldrich) according to the manufacturer’s instructions with minor modifications. 5 × 10^5^ HACs were plated in a 10 cm dish and cross-linked with final 1% formaldehyde for 10 min at room temperature. Formaldehyde crosslinking was stopped by adding glycine to a final concentration of 125mM and incubating at room temperature for 5 min. Cells were harvested and lysed to isolate nuclei in a hypotonic buffer, then re-suspended, lysed in lysis buffer, and sonicated in 1.5 ml tubes with Bioruptor Diagenode (8 × 30 s) to yield chromatin size of 100–600 bp. ChIP was performed with 2μg of anti-MEF2 (Santa Cruz sc313x) and anti-rabbit IgG. Co-precipitated DNA was then analyzed by qRT-PCR performed with SYBR green mix (Applied Biosystems). Primers sequence (FW 5’-CGTTGAGCGCTTACTGTATCT-3’; RV 5’- ACGCCCACTCCCACTATTTT-3’).

### Statistical Analysis

All the data were analysed using Prism 7 software (GraphPad), and were typically compared by a two-tailed t-test (Student`s distribution). Results are expressed as mean ± S.E.M. Probability (P) values less than 0.05 were considered to be statistically significant and have been denoted: *P < 0.05, **P < 0.01, ***P < 0.001, ****P < 0.0001.

## AUTHOR CONTRIBUTIONS STATEMENTS

SL and CM wrote the main manuscript text and prepared the figures. SL performed all experimental work with the exception of tissue harvest which CL performed. All authors reviewed the manuscript.

